# Targeting Neutrophil Extracellular Traps to inhibit Colon Cancer Tumor Necrosis and Metastasis

**DOI:** 10.1101/2025.05.30.657122

**Authors:** E. Gazzara, J. M. Adrover, A. Lui, S. Han, Z. Aminzada, N. Bhandari, N. Sivetz, V. S. Shirue, B. S. Shergill, M. B. Curtis, S. C. George, A. Cicala, A. Rishi, C. Chung, C. Devoe, H. Huang, M. Weiss, E Lou, D. A. Tuveson, S. Beyaz, P. M. K. Westcott, M. Egeblad, S. Gholami

## Abstract

Necrosis, conventionally thought of as a passive consequence of aggressive tumor growth, is associated with poor prognosis in colorectal cancer (CRC). We recently discovered that necrosis can be caused by neutrophils and neutrophil extracellular traps (NETs) aggregates driving vascular occlusion within the tumor vasculature in models of breast cancer. Here, we evaluated the role of NETs in inducing necrosis and metastasis in CRC. We found that the numbers of neutrophils primed to form NETs were elevated in the circulation of patients with CRC as compared to controls. CD177^Low^ neutrophils were also elevated, and they showed reduced extravasation capacity with intact ability to form NETs. The extent of necrosis correlated with metastasis (stage IV disease), independent of tumor size, in our human cohort. In both human and murine CRC tumors, necrotic regions were characterized by neutrophil infiltration and NET accumulation, and NET aggregates were observed in the vasculature next to the necrotic regions. Single cell RNA sequencing and spatial transcriptomic analysis of human CRC and liver metastases revealed that necrotic tumors activate pathways associated with increased metastatic potential, including epithelial-to-mesenchymal-transition. Using a mouse model of DNA mismatch repair proficient CRC, we found neutrophil infiltration and NETs increased with tumor progression. Genetic or pharmacological inhibition of NET formation decreased necrosis and metastasis, and importantly enhanced chemotherapy efficacy. Altogether, our findings show that NET formation in human CRC is a key feature of tumor necrosis, that it is associated with metastasis, and further suggest that preventing NET formation may offer clinical benefits to CRC patients.

## Introduction

Colorectal cancer (CRC) is the second most common cause of cancer-related death worldwide^1^. Over the last decade, there has been an alarming rise in incidence in patients under 50 years of age, a population that tends to develop more aggressive disease with poor survival^2^. About half of patients with CRC develop liver metastasis^3^, a major cause of mortality. While immunotherapeutic strategies have shown great benefit in various solid tumors, the majority of cases of metastatic CRC, namely those with DNA mismatch repair proficient disease, do not respond to immunotherapy and present with high recurrence rates despite liver resection and standard chemotherapy ^4^. This difference in response is related to decreased T cell infiltration and a highly immunosuppressive tumor microenvironment characterized by heavy infiltration by myeloid cells such as macrophages and neutrophils^5^. Neutrophils can exhibit a range of immunosuppressive properties, including promoting dysfunction and exclusion of cytotoxic T cells and natural killer cells, and recruiting regulatory T cells to tumors^5,6^.

Neutrophils are the most abundant leukocyte in human blood and have traditionally been described as the major drivers of acute inflammation in response to pathogenic infiltration. They have also been found to play a prominent role in cancer. Higher levels of neutrophils have been associated with poor oncologic outcomes in CRC^7–9^. Through secretion of pro-inflammatory cytokines, chemokines, and release of neutrophil extracellular traps, neutrophils contribute to tumorigenesis and progression by stimulating inflammation, such as increased production of interleukin-8^10–12^. NETs are web-like structures of extracellular DNA and cytotoxic protein granules that are released by neutrophils to capture and degrade large or supernumerary pathogens^13^. Increased levels of these highly cytotoxic structures have been found in the peripheral blood of patients with CRC, and presence of NETs in tumors may further contribute to inflammation^14^ and stimulate cancer cell proliferation^15^. Additionally, NETs play major roles in tumor progression and metastasis^16,17^, including in colorectal liver metastasis^16,18^. Some research proposes that NETs are implicated in resistance to standard therapies in many cancer types, and NET inhibition in animal models of breast cancer and CRC showed increased sensitivity to treatment^19–22^. In contrast, others have shown increased neutrophil levels to be associated with better response to chemotherapy in colorectal, gastric, and high-grade ovarian cancers. Precisely how NETs induce metastatic spread is still not fully understood and their contribution to therapeutic resistance remains unclear.

NETs have also been well described in promoting thrombosis in both benign and malignant conditions, including CRC^14^. Many studies have demonstrated the prothrombotic effects of NETs (including platelet capture, activation of coagulation factors, and clot stabilization^18,23–25^) and NET-rich thrombi have been linked to vessel occlusion and ischemic stroke in patients with underlying cancer^26–28^. While necrosis in cancer is conventionally thought to be a passive process caused by ischemia and subsequent cell death as a tumor outgrows its blood and nutrient supply, NET-induced occlusion of vessels is a plausible mechanism of tumor ischemia and subsequent necrosis. Importantly, necrosis is a poor clinical prognosticator, associated with poor disease-free and overall survival in CRC^29^. Recent studies have also shown an association between intra-tumoral necrosis and metastasis^30^, as well as between the presence of NETs and necrosis in tumors from patients with metastatic melanoma^31^. We have recently discovered that NETs deployed in the vascular luminal space drive vascular occlusion, promoting intra-tumoral hypoxia, necrosis and metastasis in breast cancer (Adrover et al., provisionally accepted). Here, we demonstrate the critical translational relevance of NET-induced necrosis and metastatic progression in primary human samples of CRC and liver metastases. DNA mismatch repair proficient CRC generally presents with high recurrence rates and has not shown any survival benefit with perioperative chemotherapy^32^. We also establish the functional relevance of NETs in driving metastasis and resistance to systemic therapy in a physiologically relevant preclinical model of DNA mismatch repair proficient CRC. These findings have important implications for translating NET-directed therapies to clinical trial implementation.

## Results

### NETs are associated with tumor necrosis in CRC patients

To explore the clinical significance of NETs in driving tumor necrosis and metastasis in CRC —we began by analyzing NETs in our prospective cohort of human CRC specimens. Notably, we found that patients with CRC exhibited significantly higher levels of circulating NETs compared to healthy volunteers (Figure 1A). Next, we analyzed tumor specimens from CRC patients using histological sections and found an abundance of NETs (defined as MPO^+^citH3^+^DAPI^+^) within these tumors (Figure 1B). NETs were predominantly localized in areas of necrosis in both primary colon tumors and liver metastases (Figure 1C and Supplementary Figure 1A-B).

**Figure 1.**
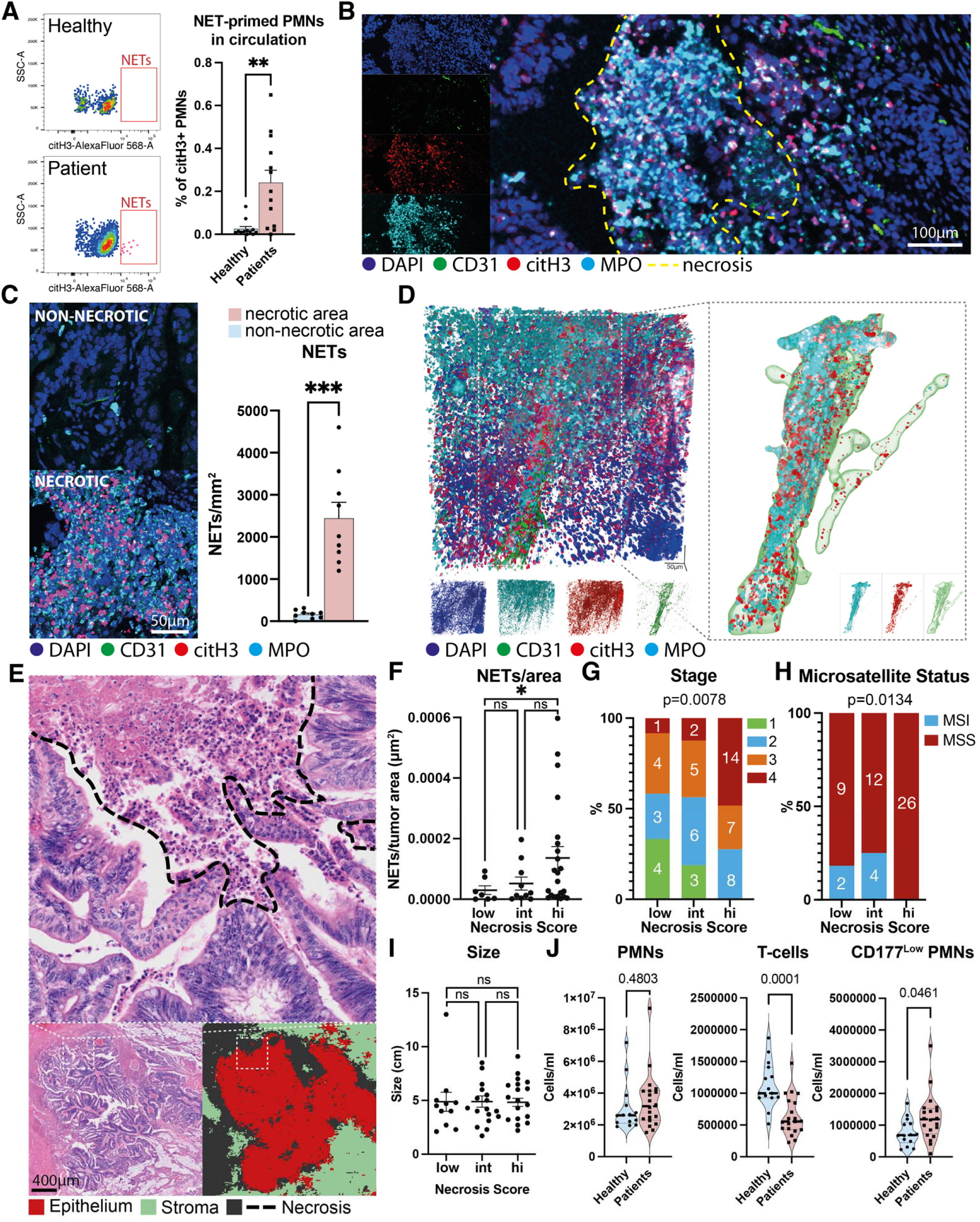
NETs are associated with necrosis in human CRC. **A)** Representative plots (left) and quantification (right) of the percentage of neutrophils poised to form NETs (CD16^+^, citH3^+^) in the blood of CRC patients compared to healthy controls, as quantified by flow cytometry. N=13 patients and N=12 healthy controls. **B)** Representative image of a human CRC tumor showing neutrophils (MPO) and NETs (MPO^+^, citH3^+^, DAPI^+^) enriched in necrotic regions (dashed yellow line). Representative of N=36. **C)** Number of NETs (MPO^+^, citH3^+^, DAPI^+^) per area of necrotic and non-necrotic regions of CRC tumors, as quantified from immunostaining. N= 9 FOV from 3 human tumors. **D)** Representative image of whole mount tissue cleared CRC tumors showing abundant intravascular NETs near necrotic regions. Representative of N=3 cleared tumors. **E)** Representative image of H&E-staining of a human CRC specimen, showing classification of tumor necrosis and stromal regions (*n* =41). **F)** Quantification of NETs per tumor area, **G)** cancer stage, **H)** microsatellite status, and **I)** tumor size in CRC tumors with no low (low), intermediate (int), and high levels of necrosis (hi). N=10 Score low, N=15 Score int, and N=16 Score hi. **J)** Quantification of neutrophils, T-cells, and CD177^Low^ neutrophils in circulation of CRC patients compared to healthy controls. N=20 patients and N=14 controls. Bars show mean ± s.e.m., **P< 0.01, ***P< 0.001, n.s., not significant, as determined by Fisher’s exact test (G, H), one-way ANOVA with Dunnett’s multiple comparison test (F, I) or unpaired two-tailed t-test (A, C, J).

Building on our prior findings that NETs cause vascular occlusion and tumor necrosis in mouse models of cancer (Adrover et al., provisionally accepted), we interrogated this mechanism in CRC using prospective analysis of freshly isolated human tumor tissue. Through whole-mount tissue clearing, we observed that not only were NETs enriched in necrotic regions of human colon tumors (Figure 1B and Supplementary Video 1), there was also significant NET accumulation within the vascular lumen of the tumor vasculature (Figure 1D, Supplementary Figure 1C, Supplementary Videos 2 and 3). Intriguingly, we also found NETs in vessels located in non-malignant colonic regions adjacent to the tumors (Supplementary Figure 1D; Supplementary Video 4), although with much lower frequency than those within the tumors. Taken together, these human data suggest that CRC induces neutrophils to form intravascular NETs primarily in the tumor vasculature, but also in the vessels of histologically normal colon.

We next assessed whether necrosis is associated with poor prognosis in our CRC patient cohort, as previously reported^29^. We quantified extent of necrosis on H&E-stained slides (Figure 1E) and stratified patients into those showing low (≤10%), intermediate (>10% and <30%), and high (≥30%) levels of necrosis (Supplementary Figure 1E-F). Tumors with high levels of necrosis had increased number of neutrophils, (Supplementary Figure 1G) and importantly, contained higher number of NETs compared to tumors with low levels of necrosis (Figure 1F). This suggests that the formation of NETs, and not merely the presence of neutrophils, is associated with necrosis. Additionally, patients who had more necrotic tumors (i.e. intermediate or high levels, compared to low levels) presented with more advanced disease (stage IV) at the time of surgery (Figure 1G), showed a higher tendency to have regional lymph node metastasis, and had an elevated ratio of neutrophils to T cells in circulation (Supplemental Figure 1H-I). Interestingly, none of the DNA mismatch repair deficient tumors, which are the only subset of CRC for which checkpoint blockade therapy is approved, had extensive necrosis (Figure 1H). This suggests that CRC patients with necrotic tumors have worse clinical prognostic characteristics compared with patients with non-necrotic tumors. Finally, we found no association between the extent of necrosis and tumor size (Figure 1I), supporting a mechanism for necrosis as an active process and not simply a secondary effect of tumors outgrowing oxygen and nutritional supply.

### CRC patients exhibit changes in circulating neutrophils

To gain a broader understanding of the neutrophil response in the systemic circulation of CRC patients and uncover a potential link between neutrophils and CRC progression, we investigated levels of circulating immune cells in our prospective cohort. We found a reduced number of T cells but no difference in total neutrophils or monocytes in circulation compared to healthy volunteers (Figure 1J and Supplementary Figure 2A). Neutrophils from CRC patients showed increased expression of adhesion molecules such as CD11b (Integrin αM) and CD11c (the fibrinogen receptor) but no differences in CD54 (ICAM-1) and CD62L (L-selectin) compared to neutrophils from healthy volunteers (Supplementary Figure 2B), potentially suggesting increased activation in circulating neutrophils in patients with CRC. Interestingly, among all circulating immune cells, the flow cytometric analysis revealed a striking change specifically in neutrophils when comparing CRC patients to healthy volunteers, emphasizing a neutrophil systemic reprogramming (Supplementary Figure 2C, note the lack of overlap between neutrophils of patients and healthy volunteers, but not in other populations). From our previous work in mouse model of breast cancer (Adrover et al., provisionally accepted), we had found Ly6G^High^/Ly6C^Low^ neutrophils were unable to extravasate in tumors but formed NETs more efficiently than classical Ly6G^High^/Ly6C^High^ neutrophils. The reduced extravasation ability of this population was critical in driving thrombo-inflammatory occlusion of the tumor vasculature and tumor necrosis. While Ly6C is not expressed in humans, we discovered that CRC patients had a greater number of neutrophils exhibiting low levels of CD177 expression compared to healthy volunteers (Figure 1J and Supplementary Figure 2C). CD177, also referred to as NB1 or PRV-1, is another member of the Ly-6 superfamily that we previously found reduced in the murine vascular-restricted Ly6G^High^/Ly6C^Low^ neutrophils, and that plays a role in the process of neutrophil extravasation^33^. We then tested whether this CD177^Low^ neutrophil population showed reduced extravasation capacity. Using an assay designed to quantify human neutrophil extravasation^34^ *in vivo*, CD177^Low^ neutrophils showed reduced extravasation ability into the human oral cavity (Supplementary Figure 2D), but they showed no defect in chemotaxis *in vitro* (Supplementary Figure 2E). Furthermore we found no differences in expression of CD10, a marker of neutrophil maturity^35^, or in nuclear morphology (Supplementary Figure 2F-G) between CD177^Low^ and CD177^Hi^ neutrophils, suggesting no difference in maturation state. Taken together, these results suggest that CD177^Low^ neutrophils, which are expanded in CRC patients, can move towards inflammatory stimuli but have reduced extravasation ability, which may result in their accumulation within the tumor vasculature.

### Cancer cells in necrotic tumors gain pro-metastatic traits

Given that necrosis within tumors is associated with poor prognosis and clinical variables associated with increased metastatic potential (Figure 1G), we next investigated how necrosis affects cancer cells in the tumor microenvironment. We performed single cell RNA-sequencing (scRNA-seq) on freshly resected liver metastases with and without necrosis from four patients undergoing liver surgery (Figure 2A and Supplementary Figure 3A). Cancer cells from these tumors separated into four distinct clusters (Figure 2B). Cluster 1 was specifically enriched in necrotic tumors (Figure 2C) and showed significantly higher expression of genes such as VIM (encodes for the cytoskeletal protein Vimentin) and SNAI1 (encodes the transcription factor Snail), among others (Figure 2D, Supplementary Figure 3B and Supplementary Table 1), suggestive of an epithelial-to-mesenchymal transition (EMT) phenotype^36,37^. VEGFA (Vascular endothelial growth factor A), which can promote angiogenesis and metastasis^38^, was also upregulated in cancer cells from cluster 1 (Figure 2D). Analysis of gene ontology (GO) terms significantly enriched within the gene signature of this cluster (Supplementary Table 2) revealed upregulation of terms related to EMT (e.g., response to TGF-β and positive regulation of EMT) and migration (e.g., negative regulation of cell-cell adhesion and positive regulation of epithelial cell migration), as well as hypoxia and the immune response such as response to IL-1, IL-8 and IL-12 neutrophil chemotaxis (Figure 2E and Supplementary Figure 3C). Of note, the signature also included a set of chemokines and cytokines known to affect neutrophil behavior^39,40^ and the hematopoietic compartment^41–43^, such as CSF1, CXCL1, CXCL2, and CXCL12, which could play a role in the systemic neutrophil changes we observed in our patient cohort (Figure 2D). Importantly several of these cytokines and chemokines, including G-CSF and CXCL2, were increased at the protein level in the plasma of CRC patients compared to healthy controls (Supplementary Figure 3D).

**Figure 2.**
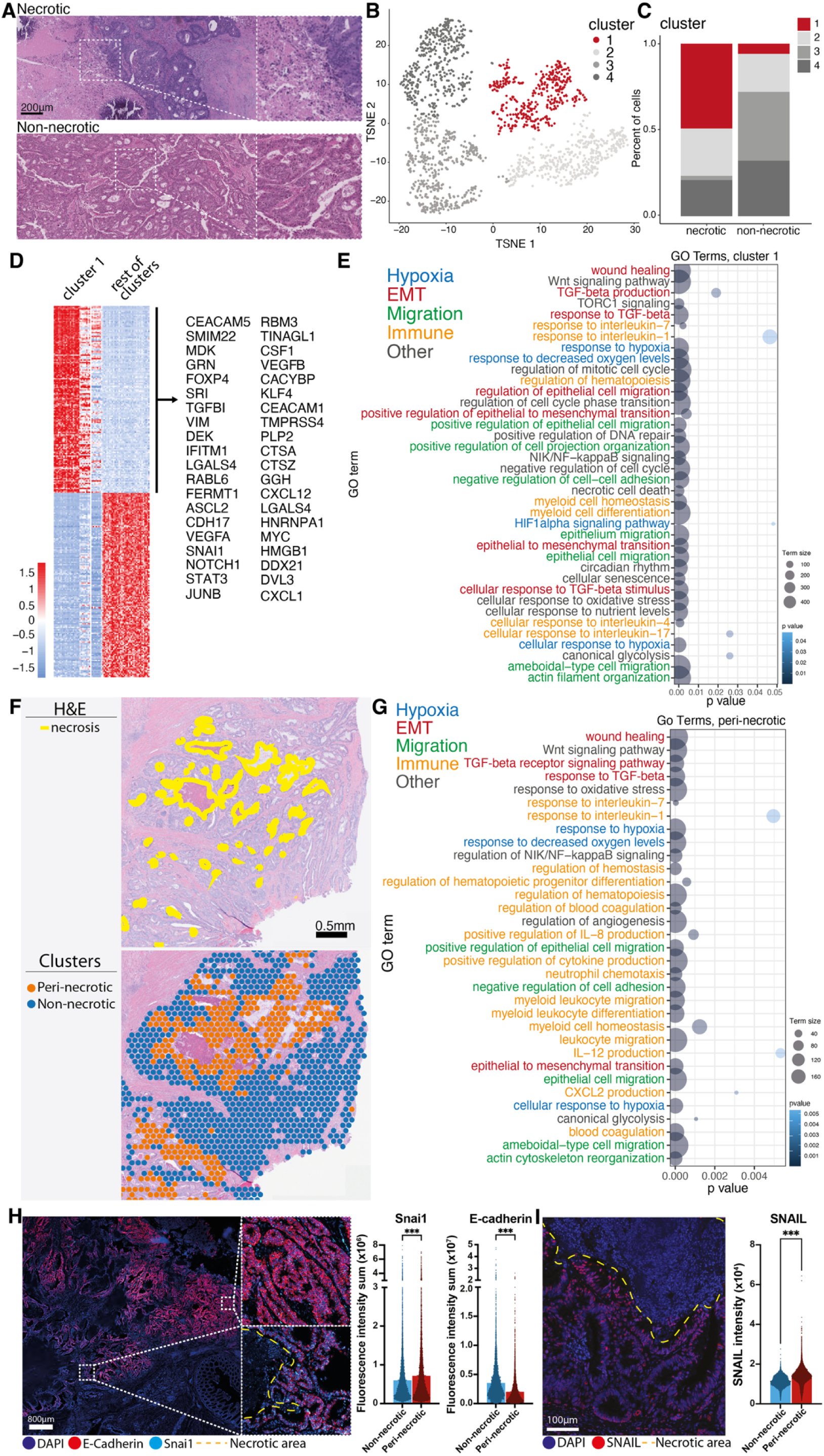
Necrotic tumors demonstrate pro-metastatic traits in human CRC. **A)** Representative images of H&E-staining of necrotic and non-necrotic human CRLM specimens used for single cell RNA sequencing. **B)** TSNE plot and **C)** quantification of epithelial cells clusters showing expansion of cluster 1 in necrotic tumors compared to non-necrotic tumors. N=2 necrotic specimens and N=2 non-necrotic specimens. **D)** Differentially expressed genes in cluster 1 as compared to clusters 2-4, selected upregulated genes shown to the right. **E)** Gene ontology enrichment analysis showing pathways related to invasion and metastasis upregulated in necrotic tumors. **F)** H&E staining (top, yellow lines delineate necrotic areas) and unbiased clustering results (bottom) in our spatial transcriptomics dataset. **G)** GO terms of the genes upregulated specifically in peri-necrotic tumor regions compared to non-peri-necrotic regions. **H)** RNAscope analysis showing increased Vimentin and decreased E-Cadherin (consistent with EMT) in peri-necrotic regions of CRC patients. **I)** IF staining of Snail in human CRC tumors, showing it increases in peri-necrotic regions at the protein level. Bars show mean ± s.e.m., ***P< 0.001 as determined by unpaired two-tailed t-test (H-I).

Necrotic tumors had more cancer cells residing in G1 phase compared to non-necrotic tumors (Supplementary Figure 3E-F), and a recent study demonstrated that TGF-β enhances the metastatic ability of cancer cells in G1 phase^44^. Furthermore, Reactome pathway analyses of the cluster 1 gene signature also revealed a significant association between tumor necrosis and TGF-β signaling, immune pathways and hematopoiesis regulation (Supplementary Figure 3G). Taken together, these data suggest that cancer cells in necrotic tumors gain pro-metastatic traits, possibly via EMT, in agreement with the clinical data from our prospective patient cohort showing increased NETs in patients with necrotic tumors and stage IV disease.

To determine the spatial localization of these cancer cells with pro-metastatic traits relative to areas of necrosis, we performed spatial transcriptomics (10x Visium ST) on two independent primary tumors from CRC patients. Unbiased clustering revealed distinct clusters corresponding to tumor regions immediately adjacent to necrosis (peri-necrotic) and regions not bordering necrosis (non-necrotic) (Figure 2F and Supplementary Figure 3H-I). Consistent with the scRNA-seq analysis (Figure 2D), we found many of the same differentially expressed genes associated with necrotic tumors downregulated or upregulated in peri-necrotic versus non-necrotic tumor regions (Supplementary Figure 3 J-K). Furthermore, genes upregulated specifically in peri-necrotic areas were enriched for pathways associated with hypoxia (e.g., response to hypoxia), EMT (e.g., epithelial to mesenchymal transition, response to TGFβ), migration (e.g., positive regulation of epithelial cell migration) and several immune-related pathways (Figure 2G). These results provide spatial context to the scRNA-seq findings (Figure 2E) and suggest that cancer cells immediately adjacent to necrosis engage in pathways that endow them with increased metastatic potential. To validate this EMT phenotype, we analyzed the RNA expression of Snail and E-cadherin by RNAscope in these tumors, which showed increased Snail and decreased E-Cadherin in peri-necrotic versus non-peri-necrotic regions (Figure 2H). Snail is a known driver of EMT, and we also confirmed its increased expression specifically in peri-necrotic regions at the protein level (Figure 2I).

### An organoid mouse model of CRC recapitulates the human phenotype

Taken together, our human data establish that NETs and tumor necrosis are strongly associated with advanced disease (stage IV) and features of malignant progression (EMT) in our patient cohort. To functionally interrogate the causal role of NETs in CRC tumor necrosis, we leveraged a model that faithfully recapitulates human CRC and metastatic disease^45–47^, using colonoscopy-guided submucosal injection of AKPS (*Apc* knockdown (*Apc^KD^); Kras^G12D^; Trp53^KO^; Smad4^KO^*) organoids into the distal colon of mice. In this model, both primary colon tumors and liver metastases present with extensive tumor necrosis, with heavy infiltration of neutrophils and NETs in necrotic regions (Figure 3A-B and Supplementary Figure 4A-B), similar to our human tissue samples. In contrast, non-necrotic tumor regions (Supplementary Figure 4C) and adjacent normal tissue regions (Supplementary Figure 4D) had few NETs, similar to the human samples (Supplementary Figure 1A-D).

**Figure 3.**
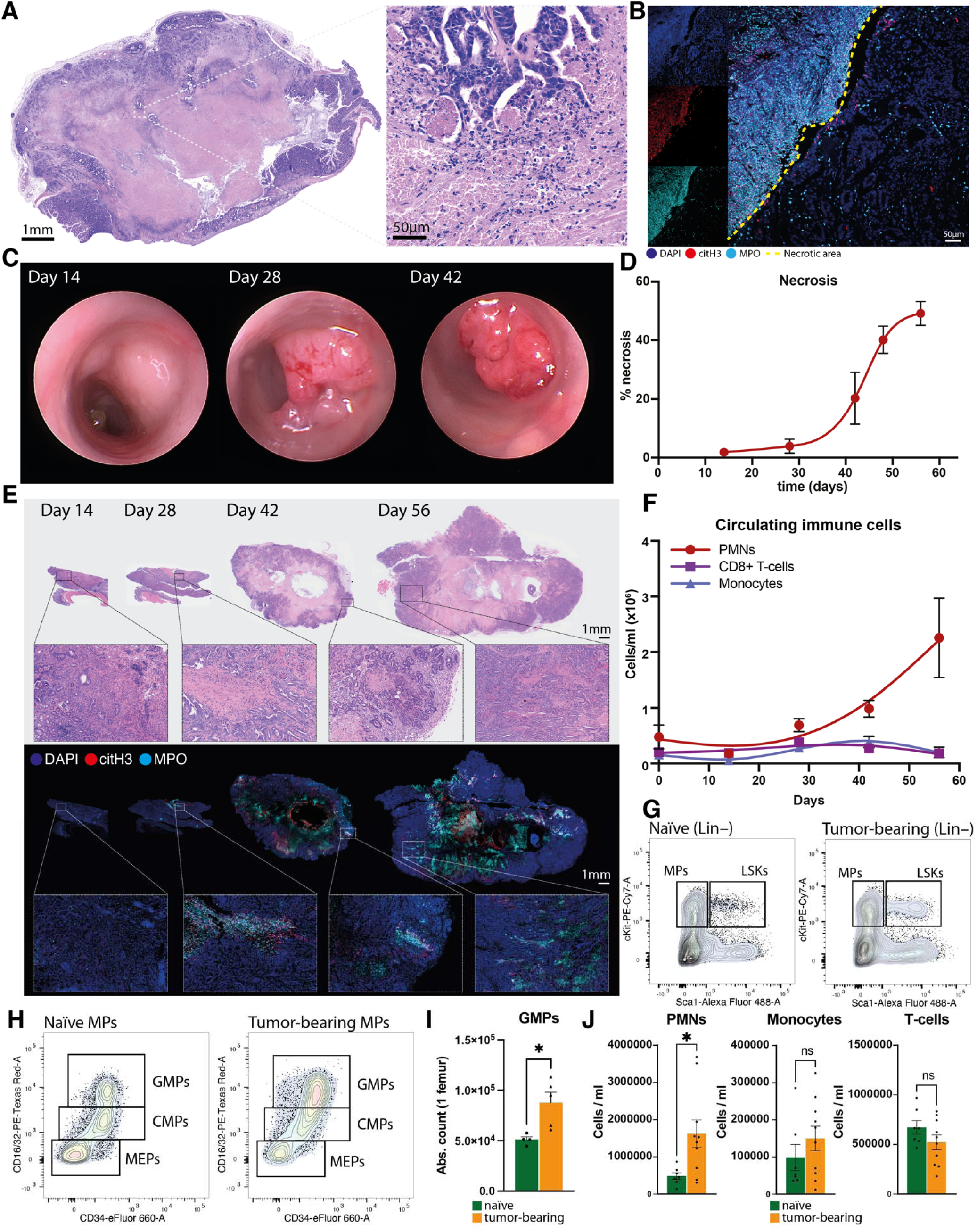
Tumor Progression Drives Systemic Neutrophil Expansion and NET Formation in Mice. **A)** Representative image of an H&E-stained AKPS-derived colon tumor, whole tumor and necrotic region (magnified). Representative of N=17 tumors. **B)** Representative image of an AKPS-derived CRC tumor showing immunostained neutrophils (MPO^+^) and NETs (MPO^+^, citH3^+^, DAPI^+^) enriched in necrotic regions (dashed yellow line). **C)** Representative images of AKPS-derived tumor development over 42 days, images obtained by colonoscopy. **D)** Quantification of percentage of necrotic tissue and **E)** representative images of AKPS tumor progression (top) and NET-formation (bottom). N=17 tumors. **F)** Quantification of immune cells in circulation of tumor-bearing mice showing increase of neutrophils during tumor development, as quantified by flow cytometry. N=3 mice per timepoint. **G)** Representative cytometry plot of lineage-negative cells in the bone marrow of tumor-bearing and naïve mice. **H)** Representative cytometry plot of myeloid progenitors (MPs) and **I)** quantification of granulocyte-monocyte progenitors (GMPs; cKit^+^, Sca1^−^, CD16/32^+^, CD34^+^) in bone marrow of tumor-bearing mice compared to naïve mice. N=5 tumor-bearing mice and N=4 naïve mice. **J)** Quantification of immune cells in circulation of tumor-bearing mice compared to naïve mice. N=10 tumor-bearing mice and N=7 naïve mice. Bars show mean ± s.e.m., *P< 0.05, n.s., not significant, as determined by unpaired two-tailed t-test.

Having confirmed that this model recapitulates the necrosis phenotype observed in human CRC, we assessed the extent of necrosis throughout an eight-week observation period of CRC progression (Figure 3C). AKPS tumors exhibited extensive neutrophil- and NET-rich necrotic regions (Figure 3D-E) as early as 28 days after implantation. Notably, tumor size at day 28 was still small, suggesting that necrosis is not solely a feature of late-stage and large tumors. Spontaneous metastases were first apparent at day 42 (1.54% of total liver area) and extensive by day 48 (19.62% of total liver area) after tumor implantation (Supplementary Figure 4E). Metastatic progression was associated with increased circulating neutrophils, but not cytotoxic T cells or monocytes (Figure 3F), which mirrored an increase of infiltrating neutrophils in the primary colon tumors and liver metastases (Supplementary Figure 4F).

Our human scRNA-seq analysis revealed increased expression of genes related to myelopoiesis in necrotic tumors (Figure 2D and Supplementary Figure 3G). We next examined the bone marrow compartment of tumor-bearing mice at day 56 after implantation. We found that relative to naïve mice, tumor-bearing mice harbored increased numbers of early hematopoietic stem (LSKs: lineage negative, Sca-1 positive, c-Kit positive) and myeloid progenitor cells in the bone marrow (Figure 3G and Supplementary Figure 4G-H). Interestingly, tumor-bearing mice had increased numbers of granulocyte-monocyte progenitors (GMPs) in the bone marrow (Figure 3H-I) and an expansion of mature neutrophils in circulation, but not monocytes or T cells (Figure 3J). These data show that the primary colon tumors induce a myeloid skew in the hematopoietic compartment, enhancing the production of circulating neutrophils that likely drive the observed increase in tumor neutrophil infiltration, NETs, and necrosis.

### Genetic and pharmacologic NET inhibition reduce tumor necrosis and metastasis

To functionally interrogate the causal relationship between NETs and tumors necrosis, we implanted AKPS tumors in neutrophil-specific Protein Arginine Deaminase 4 (PAD4) deficient mice (Mrp8^Cre^; Padi4^fl/fl^; hereafter referred to as PAD4^ΔN^), that do not form citrullinated NETs (particularly citrullinated histone 3-positive NETs—which are formed during a process requiring PAD4). As expected, tumor-bearing PAD4^ΔN^ mice had a lack of MPO^+^citH3^+^DAPI^+^ NETs within tumors but the number of circulating immune cells was the same as in tumor-bearing Mrp8^Cre^-negative Padi4^fl/fl^ littermates (PAD4^WT^) (Supplementary Figure 5A-B). Importantly, while they showed no difference in primary tumor weight (Supplementary Figure 5C), PAD4^ΔN^ mice had significantly reduced necrosis in AKPS colon tumors (Supplementary Figure 5D), demonstrating that NETs are key drivers of necrosis in CRC. In agreement with our clinical observations that necrosis is associated with higher stage disease and metastasis, the decreased necrosis in PAD4^ΔN^ mice was accompanied by significantly reduced liver metastases (Supplementary Figure 5E). This suggests that NET-induced necrosis drives cancer cells to gain pro-metastatic traits and the ability to disseminate from the primary tumor more efficiently, as predicted by our human scRNA-seq analysis (Figure 2E). Alternatively, NETs could play a role in recruiting or supporting the survival of early metastatic cells in the liver. To interrogate this alternative hypothesis, we developed an experimental metastasis model by injecting AKPS cells directly into hepatic circulation of PAD4^WT^ and PAD4^ΔN^ mice via the portal vein. In this model, we observed no differences in liver engraftment between mice proficient and deficient in NET formation (Supplementary Figure 5F), suggesting that the enhanced metastatic progression driven by NET-induced necrosis is likely due to increased cancer cell shedding from the primary tumor.

While deletion of the gene encoding for PAD4 in neutrophils is a valuable tool to study the role of NETs in the preclinical setting, it is not clinically tractable. Thus, we sought to explore the potential of pharmacologic NET inhibition to reduce necrosis and metastasis in our CRC model. We and others recently showed that disulfiram, an FDA-approved drug, can efficiently reduce NET formation in human and rodent neutrophils^48–50^. AKPS tumor-bearing mice treated with disulfiram showed significantly fewer NETs (Figure 4A-B) and reduced necrosis in tumors (Figure 4C) compared to control mice. Although disulfiram has diverse effects beyond inhibition of NET formation^51^, these results further support that NETs drive necrosis formation. As with genetic targeting of NET formation, we found no difference in tumor size between disulfiram and vehicle treated mice (Figure 4D), and only minor differences in composition of circulating immune cells (Supplementary Figure 5G). Finally, disulfiram treatment significantly reduced the rate of liver metastases compared to control mice (Figure 4E). Together with the reduction of primary tumor necrosis, this further suggests that NET-driven necrosis within colon tumors promotes metastatic spread.

**Figure 4.**
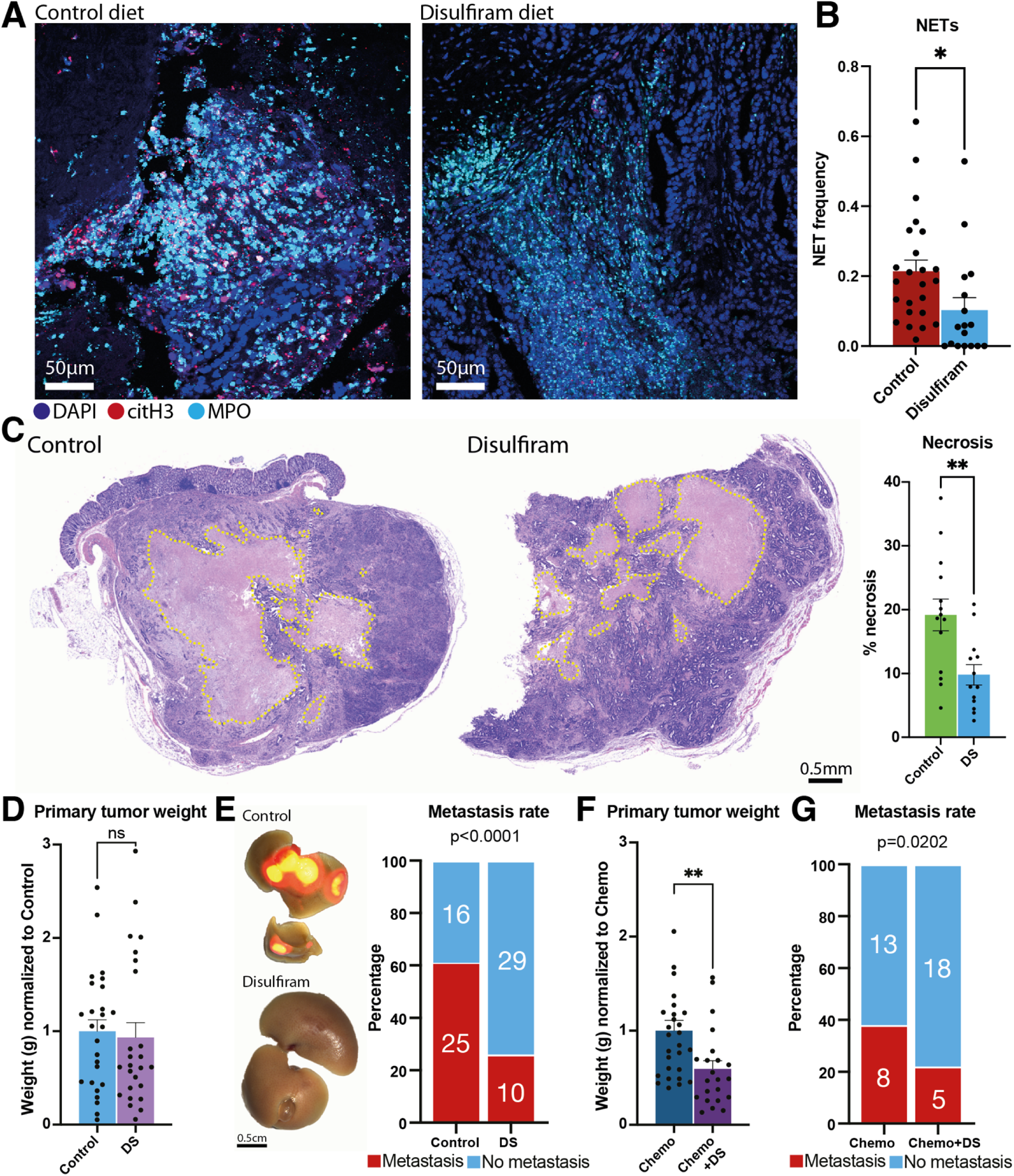
NET-inhibition reduces tumor necrosis, size, and metastasis and enhances response to chemotherapy. **A)** Representative immunostaining and **B)** quantification of NETs in colon tumors of control and disulfiram-treated mice. N=24 control mice and N=17 disulfiram-treated mice (DS). **C)** Representative H&E-stained images and quantification of necrosis within the colon tumors of mice treated with disulfiram compared to control mice. N=13 disulfiram-treated mice and N=14 control mice. **D)** Tumor weight at endpoint of disulfiram-treated mice normalized to control mice. N=26 controls and 25 disulfiram-treated mice. **E)** Representative brightfield images with overlying tumoral mScarlet fluorescence and quantification of the rate of metastasis in AKPS-bearing mice treated with disulfiram compared to control mice. N=10 control mice and N=7 disulfiram-treated mice. **F)** Primary tumor weight at endpoint of mice treated with chemotherapy and with combination of disulfiram with chemotherapy, normalized to chemotherapy group. Chemotherapy-treated (Chemo; N=17), combination therapy-treated (Chemo+DS; N=17). **G)** Quantification of rate of liver metastasis in AKPS-bearing mice treated with chemotherapy alone or disulfiram together with chemotherapy. Bars show mean ± s.e.m., *P< 0.05, **P< 0.01, n.s., not significant, as determined by unpaired two-tailed t-test (B, C, F), Fisher’s exact test (E, G) or Mann-Whitney test (D).

### Pharmacologic NET inhibition and necrosis reduction enhance CRC standard of care therapy

Given our previous results in breast tumors, which revealed hypoperfusion in necrotic regions and peri-necrotic regions as a result of NET infiltration (Adrover et al., provisionally accepted), and our demonstration that CRC tumors have similar areas of vessel occlusion (Figure 1D, Supplementary Figure 1C and Supplementary Video 3), we hypothesized that reduction of vascular occlusion and necrosis, through NET blockade, could enhance the bioavailability of systemic therapy to the tumor. To test this hypothesis, we treated AKPS tumor-bearing mice with systemic standard-of-care chemotherapy for CRC (FOLFOX: 37.5mg/kg 5-fluorouracil and 6mg/kg oxaliplatin) with or without concurrent administration of disulfiram. Mice treated with the combination of chemotherapy and disulfiram showed a significant reduction in primary colon tumor size compared to mice treated with chemotherapy alone (Figure 4F and Supplementary Figure 5H). Combination chemotherapy and disulfiram treatment also reduced metastatic rate (Figure 4G) compared to chemotherapy alone. The combination treatment was well-tolerated (Supplementary Figure 5I) and lowered the number of neutrophils and monocytes in circulation relative to mice treated with chemotherapy alone (Supplementary Figure 5J). These data establish that decreased tumor necrosis, achieved by blocking NETs, can augment the effects of systemic chemotherapy in primary tumors and suppress liver metastasis compared to standard-of-care chemotherapy.

## Discussion

Our findings demonstrate that NETs drive tumor necrosis and metastasis in CRC, highlighting a distinct reprogramming with strong human translational relevance and therapeutic potential in CRC progression. CRC poses a significant challenge due to frequent liver metastases and high recurrence rates in the liver despite surgical resection (over 70%). This is particularly critical in DNA mismatch repair proficient tumors, which do not show any survival benefit from peri-operative and adjuvant chemotherapy, underscoring the urgent need for better therapeutic strategies.

Our findings support a model in which CRC tumors influence the hematopoietic compartment to drive a myeloid skew, resulting in enhanced production of neutrophils in the circulation (Figure 3H-J). These neutrophils in turn drive necrosis within primary tumors, creating peri-necrotic regions where cancer cells acquire pro-metastatic traits, ultimately fueling metastasis to the liver. This model is strongly supported by our patient cohort data and aligns with previous literature showing that tumor necrosis correlates with worse prognosis in CRC^29^ and other cancers^52–55^. Using a comprehensive, multi-platform approach—including scRNA-seq, 3D imaging, and spatial transcriptomics—we demonstrate that NETs aggregate in the tumor vasculature, that they accumulate in necrotic regions of both primary tumors and liver metastases, and that the cancer cells and microenvironment surrounding the NET-associated necrotic regions is reprogrammed towards EMT and metastatic progression (Figure 2D-G). Moreover, we show that targeting NETs not only reduces tumor necrosis and metastasis (Figure 4C-E) but also enhances the efficacy of chemotherapy (Figure 4F-G) in a highly relevant mouse model that recapitulates the histopathological progression (Figure 3A-F) and immunotherapy refractory nature of human DNA mismatch repair proficient CRC^47^. These findings provide a compelling rationale for integrating NET-targeting strategies into future clinical trials for DNA mismatch repair proficient CRC patients, offering a novel therapeutic avenue to improve patient outcomes.

NETs have been implicated in most steps of tumor progression^17^, including in CRC^56^. Therefore, as an initial step of investigation, we utilized our human samples to understand the role of NETs in CRC. Interestingly, we found circulating neutrophils with citrullinated histone 3, indicative of nuclear decondensation and NET formation, despite these neutrophils being viable and intact (Figure 1A). This suggests that neutrophils in the early stage of NET formation are present in the circulation of cancer patients, concordant with our findings of NETs in extra-tumoral tissues such as the peritumoral colon (Supplementary Figure 1D). We propose a model in which neutrophils form NETs in the vasculature of tumors, leading to thromboinflammatory occlusion of the vessels, subsequent hypoxia and necrosis of the regions downstream of that vascular bed. NETs are indeed highly pro-thrombotic structures, and neutrophil-platelet interactions are drivers of NET formation^23,28^. Interestingly, in our cancer patient cohort we found an expansion of CD177^Low^ neutrophils, which have lower extravasation ability (Supplementary Figure 2D) therefore are likely to accumulate intraluminally and contribute to thromboinflammatory occlusion of the vessels.

In our patient specimens, we found neutrophil aggregates and NETs inside the vasculature (Figure 1D and Supplementary Figure 1C) as well as in necrotic areas (Figure 1C and Supplementary Figure 1A-C). It is interesting to note that we did not find a correlation of tumor necrosis with tumor size in our patient cohort (Figure 1I), which challenges the conventional idea that necrosis is a passive process resulting from tumor growth outpacing blood supply. We also found that the presence or absence of necrosis within liver metastases in our preclinical model was not linked with metastatic foci size. That is, in the same liver section, some metastatic lesions were necrotic while others were not (Supplementary Figure 4A), again suggesting that the formation of necrosis is not related to the size of the lesion. This finding is congruent with our proposed model, as the initial step of neutrophil aggregation in the vasculature and NET formation leading to vascular occlusion may be a stochastic event in some of the lesions, regardless of tumor size.

As tumor necrosis has been well described to be associated with poor prognosis and metastasis in CRC patients, we sought to understand the underlying mechanism by which necrosis enhances metastatic spread. We first generated scRNA-seq data of necrotic vs. non-necrotic tumors to understand transcriptomic differences of cancer cells in the presence and absence of necrosis. We found that necrotic tumors demonstrated an enrichment of a particular cluster of cancer cells that showed a gene expression profile consistent with the initiation of EMT (Figure 2D-E and Supplementary Figure 3B-C, E-G), including increased expression of *VIM* (a mesenchymal marker) and *SNAI1* (Snail, a well-known driver of EMT^57,58^) expression. We also observed enrichment of pathways related to hypoxia (a known inducer of EMT^59^) and epithelial cell migration (Figure 2E). Given these observations and the fact that necrotic regions showed areas of vessel occlusion (Supplementary Video 3), we sought to understand whether cancer cells immediately adjacent to necrotic areas are in a hypoxic state and express EMT/pro-metastatic traits. To test this hypothesis, we performed spatial transcriptomic analyses of primary tumors from CRC patients. We found that cancer cells in the vicinity of necrosis (peri-necrotic regions) showed a similar pattern of up- and down-regulated genes as the cluster we found enriched in necrotic tumors by scRNA-seq (Supplementary Figure 3J-K), suggesting that indeed the changes we found in our scRNA-seq were taking place specifically in peri-necrotic regions of these tumors. Importantly, when we interrogated the pathways changing specifically in peri-necrotic regions compared to non-peri-necrotic regions of those tumors, we found very similar pathways encompassing EMT, hypoxia and migration (Figure 2G). As a tumor becomes more necrotic, more tumor cells are exposed to the necrotic microenvironment at the peri-necrotic border, increasing the number of cancer cells that may acquire the pro-metastatic traits of EMT and invasion. This results in a higher metastatic potential for these tumors, a conclusion that is consistent with our data from human CRC and clinical reports that connect necrosis with metastasis^29^.

To understand the role of NETs in tumor necrosis, we turned to our colonoscopy-guided, AKPS preclinical model. We first confirmed that the preclinical model closely recapitulated critical features of human CRC (Figure 3A-F). Importantly, when we genetically (using PAD4^ΔN^ mice) or pharmacologically (using disulfiram) reduced the ability of neutrophils to form NETs, we found reduced necrosis in primary tumors (Supplementary Figure 5D and Figure 4C), demonstrating that NETs are indeed key drivers of tumor necrosis in CRC. Of note, tumor size was not different between our NET-deficient models and their respective controls (Supplementary Figure 5C and Figure 4D). This again argues that necrosis is not a passive phenomenon secondary to tumor growth, but an active phenomenon driven by neutrophils and NETs. In agreement with our hypothesis that necrosis drives metastasis, blocking NETs led to reduced metastatic burden in the liver (Supplementary Figure 5E and Figure 4E). This is in line with recent reports using an orthogonal NET targeting approach with DNase I^60^. But it is important to note that NETs have been shown to be critical for the homing of cancer cells to the liver^12^, so it was possible that our reduced metastatic spread was due to reduced uptake in the liver rather than to reduced shedding from the primary tumor. To test this hypothesis, we used an experimental metastasis model where we bypass the shedding from the primary tumor and test the homing ability of cancer cells in the presence and absence of NETs. In this case, the differences observed in the liver metastasis burden were lost (Supplementary Figure 5F). This suggests that the reduction in metastatic burden in the liver observed in our NET-deficient model is driven by decreased shedding of cancer cells from the primary tumor, rather than homing of the cells to the liver.

Finally, because necrotic regions are avascular (Supplementary Video 1) and adjacent peri-necrotic regions are hypoperfused, we hypothesized that NET inhibition could be beneficial in combination with standard of care chemotherapy. We found that the combination treatment with disulfiram and chemotherapy showed enhanced responses both in the primary tumor as well as in liver metastases (Figure 4F-G), which agrees with previous literature suggesting that the presence of NETs within tumors may hinder the efficacy of chemotherapy^61^. We acknowledge however, that disulfiram has pleiotropic effects^62^ beyond reducing NET-formation. Although we had more specific means to block NET formation (the genetic PAD4^ΔN^ model), we used disulfiram in these set of experiments as disulfiram is an FDA-approved drug which could be translated into future human clinical trials. Given the effects of disulfiram in decreasing metastases, it could be indicated in the adjuvant setting when treating primary CRC tumors. Because disulfiram exhibits effects beyond NET inhibition, future research should focus on identifying more selective agents that either block NET formation or promote their clearance, enabling more targeted combination therapies.

In summary, the results of this study demonstrate that colorectal tumor necrosis is not a passive phenomenon, but an active process driven by NETs, which can be clinically targeted to reduce metastatic spread and potentially improve patient survival. Of particular translational relevance is our finding that combination treatment with NET inhibitors and standard of care chemotherapy improved response compared to chemotherapy alone. Future studies will be essential to validate these findings in clinical settings and to develop more specific NET-targeting agents for integration into combination therapies.

## Acknowledgements

This work was supported by funding provided to M.E. by the National Institutes of Health (NIH 1R01CA2374135), The Giovanis Institute for Translational Cell Biology, and the International Neutrophil Institute. J.M.A. was the recipient of a Cancer Research Institute Irvington Fellowship (CRI Award #3435). S.G. was awarded funding through CSHL/Northwell Health (Award # 60430598). P.M.K.W and members of his laboratory were supported by NCI support grant P30-CA045508, the V Foundation for Cancer Research (V Scholar Award), the Betty Ajces Trust, the Wasily Family Foundation, and the Don Monti Memorial Research Foundation. N.S. was supported by an NIH award (NIH 1F31CA281246-01). This work was performed with assistance from Cold Spring Harbor Laboratory Shared Resources, which are supported by Cancer Center Support Grant P30CA045508. We thank Shared Resource heads Drs. Jonathan Preall, Rachel Rubino, Pamela Moody, and Qing Gao for guidance and assistance. We also thank the Northwell Health Biorepository for assisting in patient identification and human sample collection.

## Materials and methods

### Human samples

All human specimens were collected by Northwell Health Biorepository (NHBR). Patients over 18 years of age with a diagnosis of primary colorectal cancer with or without metastasis to distant organs were included. We excluded patients with prior or ongoing infection with pathogens such as Hepatitis B virus (HBV), Hepatitis C virus (HCV 3.0) and Human Immunodeficiency viruses, Types 1 and 2 (HIV 1,2). Archival FFPE tissue samples from the same patient population were also provided by NHBR. Clinical data was obtained from Northwell Health electronic medical record. This study was approved by the Institutional Review Board and NHBR (Protocol #1810 and UCD IRB protocol #24-0635) at Feinstein Institutes for Medical Research Northwell Health.

### Mice

All experiments were performed in 6- to 15-week-old mice of approximately equal genders, housed in specific pathogen-free facilities at Cold Spring Harbor Laboratory (CSHL), with food and water available ad libitum and in a 12h:12h light/dark regime. C57BL/6 mice were obtained from The Jackson Laboratory. Pad4^ΔN^ mice were generated in-house by crossing Mrp8^CRE^ mice with Padi4^fl/fl^ mice and were compared to Cre-negative littermates. All mice were acclimated to the animal housing facility at least for one week prior to experimentation.

All procedures were approved by the CSHL Institutional Animal Care and Use Committee and were conducted in accordance with the NIH’s *Guide for the Care and Use of Laboratory Animals*.

### Organoid culture

mScarlet-positive AKPS organoids were maintained in 20uL domes of 80% Matrigel in A+++ media (1% Penicillin-Streptomycin (Thermo Fisher), 1% glutamax (Thermo Fisher), and 2% B27 (CSHL)). Organoid cultures were thawed independently for every experiment to avoid genetic drift, and passaged every 3 days. TrypLE Express Enzyme (Thermo Fisher, no phenol red) was used to dissociate organoids.

### Tumor models

AKPS organoids from day-3 culture were harvested and washed with cold phosphate buffered saline (PBS) to melt Matrigel domes. Organoids were injected into the submucosa of the distal colon via colonoscopic guidance (1x10^6^ organoids in 100uL of 20% Matrigel in OptiMEM (Thermo Fischer)), as previously described.^63^ Tumor engraftment was confirmed through colonoscopy 2 weeks after injection. Portal vein injections were performed using day-2 culture AKPS organoids digested into a single cell solution using TrypLE for 12 minutes. 1x10^6^ cells were injected in 100uL OptiMEM into the portal vein.

### NET inhibition with disulfiram

Gamma-irradiated Purina rodent chow diet #5053 repelleted (as control) and Purina chow diet #5053 with 1g of disulfiram per kg of diet (experimental) were purchased from Research Diets, Inc. Mice were fed with the control or experimental diet ad libitum starting prior to and for the duration of the experiment. Inhibition of NET formation was previously reported in a model of acute lung injury^48^, and was confirmed here in all the experimental groups.

### Treatment with standard of care chemotherapy

Treatment began 2-3 weeks following organoid injection, after tumor engraftment in the colon was confirmed via colonoscopy. Mice were treated with 6mg/kg Oxaliplatin (Sigma, #O9512-5MG) and 37.5mg/kg 5-fluorouracil (Sigma, #F6627-5G) or vehicle (3.75% dimethyl sulfoxide in PBS) once a week for 5 weeks via intraperitoneal injection. All mice were sacrificed 9-11 weeks post-organoid injection. Mice that were treated with combination chemotherapy and disulfiram were maintained on a disulfiram diet for the duration of the experiment. Mice were monitored for adverse effects such as weight loss and deconditioning for the duration of treatment. Humane endpoints for sacrifice (weight loss greater than 20%, unrecoverable deconditioning, obstruction from colonic tumor) were used for all experiments.

### Analysis of tumor necrosis in CRC and CRLM

Tissues were sectioned at the CSHL Animal & Tissue Imaging shared resource or by the NHBR. Tissue slides were stained with hematoxylin & eosin. The necrotic area was calculated using QuPath’s (v0 4.4) MLP-ANN pixel classifier trained to the necrotic regions and compared to total tumor area (excluding adipose tissue, also trained in the classifier). Slides were scored for percent area of necrosis per area of entire tumor. Patients were classified into groups by percent necrosis: low necrosis (<10%), intermediate necrosis (<30%), and high necrosis (≥30%).

### Quantification of metastatic burden in mouse liver and lungs

Mice were sacrificed between 8-11 weeks post AKPS injection. Lung and liver tissue were immediately excised and submerged in 10% Neutral Buffered Formalin (NBF) and fixed at room temperature for 24 hours. Tissue was imaged with Leica M205c stereoscope. Metastatic lesions were identified by presence of endogenous *m*Scarlet fluorescence in tissue. FIJI (ImageJ) was used to determine the area of each metastatic lesion. Metastatic burden was quantified as metastatic area/total tissue area and was summed for each group.

### Immunofluorescence imaging

To stain formalin-fixed paraffin-embedded (FFPE) slides, paraffin sections were first deparaffinized and rehydrated, and antigen retrieval was carried out by boiling slides in Tris EDTA buffer (10 mM Tris Base, 1 mM EDTA, pH 9.0) for 10 minutes in a pressure cooker. Tissues were incubated with a blocking buffer containing PBS with 25% FBS, 0.1% Triton X-100 and 0.2% BSA for 30 minutes at room temperature. The tissues were incubated with the corresponding primary antibodies (see “Antibodies for immunofluorescence staining table” below) overnight at 4°C. The next day, the samples were washed three times in PBS and stained with the corresponding secondary antibodies for 2 hours at room temperature. Then, the samples were washed, counterstained with DAPI, and mounted with antifade medium (ProLong Gold, Thermo Fisher, cat #P10144).

### Whole mount immunostaining and tissue clearing

Mice were euthanized with CO_2_, and their tissues were immediately excised and submerged in 10% NBF and fixed at room temperature for 24 hours. Human tissue was fixed similarly upon receipt. After three washes with PBS for 1 hour each at room temperature, the tissues were permeabilized in methanol (MetOH) gradients in PBS (PBS > MetOH 50% > MetOH 80% > MetOH 100%, for 30 minutes in each solution). Then, the tissues were bleached with Dent’s bleach (15% H_2_O_2_, 16.7% dimethyl sulfoxide [DMSO] in MetOH) for 1 hour at room temperature and rehydrated through descending methanol gradients in PBS (MetOH 80% > MetOH 50% > PBS, 30 minutes in each solution). Then, the tissues were incubated with a blocking buffer containing PBS with 0.3% Triton X- 100, 0.2% BSA, 5% DMSO, 0.1% azide, and 25% FBS overnight at 4°C with shaking. Afterwards, the tissues were stained with primary antibodies (see “Antibodies for immunofluorescence staining table” below) at 1:200 in blocking buffer for 3 days at 4°C with shaking. After washing for 24 hours in washing buffer (PBS with 0.2% Triton X-100 and 3% NaCl), the tissues were stained with secondary antibodies (see “Antibodies for immunofluorescence staining table” below) at 1:400 for 3 days at 4°C with shaking. Then, the tissues were washed for 24 hours in washing buffer and thereafter dehydrated in MetOH gradients in dH_2_0 using glass containers (MetOH 50% > MetOH 70% > MetOH 90% > 3x MetOH 100%, 30 minutes for each step). The tissues were then cleared for 30 minutes in 50% MetOH and 50% benzyl alcohol, benzyl benzoate (BABB, mixed 1:2) and for 1 hour in 100% BABB, and finally, imaged on an SP8 microscope (Leica, typical Z-depths of 300–500 µm). Quantification was performed with Imaris software (Bitplane).

### Immunofluorescence image quantification

NETs in tissue were quantified in Bitplane Imaris. Briefly, the triple colocalization of DAPI, citH3, and MPO was used as the criterion to detect and quantify the NETs using Imaris spots.

### Antibodies for immunofluorescence staining

| *Antibody against* | *Clone* | *Manufacturer* | *Catalogue #* |
| --- | --- | --- | --- |
| *citH3* | Polyclonal | Abcam | Ab5103 |
| *MPO* | Polyclonal | R&D Systems | AF3667 |
| *CD31* | MEC 13.3 | BD Biosciences | 55027 |
| *CD3* | 17A2 | BioLegend | 100202 |
| *SNAIL* | Polyclonal | Abcam | ab85936 |

### Secondary antibodies

| *Antibody against* | *Manufacturer* | *Catalogue #* |
| --- | --- | --- |
| *Donkey-anti-rabbit-AF568* | Invitrogen | A10042 |
| *Donkey-anti-rat-AF488* | Invitrogen | A212008 |
| *Donkey-anti-goat-AF647* | Invitrogen | A21447 |
| *Donkey-anti-rabbit-Dy594* | Thermo Fisher | SA5-10040 |

### Flow cytometry and cell sorting

Flow cytometric analyses were performed using a Fortessa Analyzer (BD Biosciences). The analysis was performed using FlowJo v10 (Tree Star Inc.). Cell sorting experiments were performed using a fluorescence-activated cell sorting (FACS) SONY cell sorter (SONY Biotechnology). All analyses were conducted at the Flow Cytometry Shared Resource at CSHL. Absolute quantification was done using Trucount absolute counting beads (BD Biosciences, #340334) according to the manufacturer’s instructions.

Neutrophils were isolated from blood by FACS. Briefly, blood was drawn into EDTA-coated tubes, red blood cells were lysed in ACK (Ammonium-Chloride-Potassium, Gibco, USA, #A1049201) lysis buffer (Thermo Fisher, USA), and blood cells were stained with antibodies against CD16 and CD177 (see antibody table below, immune panel). Immediately prior to flow cytometric analysis, DAPI was added to the cells. Then DAPI-, CD16+ neutrophils were sorted. From this neutrophil population, CD177-HI and CD177-LOW neutrophils were sorted.

Cytometric analyses of blood and tissues from naïve, tumor-bearing mice, CRC patients, and healthy volunteers were performed as previously reported^64^. Briefly, tissues (tumor, liver, lung, and bone marrow) were extracted, kept in cold PBS (except liver, kept at room temperature in HBSS), and processed immediately. Tissues were digested in Hanks’ balanced salt solution (HBSS) with collagenase (1 U/ml, Cell Technologies) and DNase I (1 mU/ml, Sigma) for 30 minutes at 37°C. Bone marrow was mechanically dissociated to prepare single-cell suspensions by flushing and straining, respectively. Leukocytes in liver were enriched by centrifugation at 800 x g for 30 minutes, using a 36% Percoll (GE Healthcare, diluted in HBSS) gradient and red blood cells were lysed in ACK hypotonic buffer. Single-cell suspensions from all tissues were incubated with fluorescently-conjugated antibodies using the hematopoietic panel below (bone marrow) or the immune panel (all tissues).

### Hematopoietic panel

| *Antibody against* | *Clone* | *Manufacturer* | *Catalogue #* |
| --- | --- | --- | --- |
| *CD34 eFluor 660* | RAM34 | Thermo Fisher | 50-0341-82 |
| *CD48 APC/Cy7* | HM48-1 | BioLegend | 103432 |
| *CD117 (cKIT) PE/Cy7* | 2B8 | BD Biosciences | 558163 |
| *CD150 (SLAM) BV510* | TC15-12F12.2 | BioLegend | 115929 |
| *Ly6A/E (Sca-1) FITC* | D7 | Thermo Fisher | 11-5981-82 |
| *CD16/32 PE* | 93 | Thermo Fisher | 12-0161-82 |
| *Biotin mouse lineage* | Cocktail | BD Biosciences | 558451 |
| *Streptavidin DyLight 405* | N/A | Jackson Laboratory | 016-470-084 |

### Mouse immune panel

| *Antibody against* | *Clone* | *Manufacturer* | *Catalogue #* |
| --- | --- | --- | --- |
| *Ly6G-AF647* | 1A8 | BioLegend | 127610 |
| *CD3-AF488* | 17A2 | BioLegend | 100210 |
| *CD45-BV785* | 30-F11 | BioLegend | 103149 |
| *CD11b-PE* | M1/70 | BioLegend | 101208 |
| *CD49b-APC/Cy7* | M5/114.15.2 | BioLegend | 108920 |
| *CD8a-PcP/Cy5.5* | SA044G4 | BioLegend | 100734 |
| *Ly6C-PE/Cy7* | HK1.4 | BioLegend | 128018 |
| *CD11c-BV650* | N418 | BioLegend | 117339 |

### Human immune panel

| *Antibody against* | *Clone* | *Manufacturer* | *Catalogue #* |
| --- | --- | --- | --- |
| *CD56-AF488* | 5.1H11 | BioLegend | 362518 |
| *CD45-BV785* | HI30 | BioLegend | 304048 |
| *CD11b-PE* | ICRF44 | BioLegend | 982606 |
| *CD177-APC/Cy7* | MEM-166 | BioLegend | 315810 |
| *CD55-PcP/Cy5.5* | JS11 | BioLegend | 311315 |
| *CD10-PE/Cy7* | BB27 | BioLegend | 331014 |
| *CD14-BV650* | M5E2 | BioLegend | 301836 |
| *CD16-APC* | 3G8 | BioLegend | 301836 |
| *CD3-BV421* | OKT3 | BioLegend | 317344 |

### NET-primed neutrophils in circulation

100uL of whole blood was lysed and stained with antibodies against CD45, CD14, CD16, and CD177 (see antibody table below, immune panel). Cells were then fixed and permeabilized using Cyto-Fast^TM^ Fix/Perm Solution kit (Biolegend) for 20 minutes at 37°C in the dark. Intracellular citrullinated histone 3 was stained for using anti-citH3 (Abcam, # Ab5103) 1:200 in wash buffer for 15 minutes at 4°C. Cells were washed and stained using secondary antibody donkey-anti-rabbit-AF568 (DARB568, Invitrogen, #A10042) 1:500 for 15 minutes at 4°C. A secondary only sample was stained for cell surface markers and DARB568 only. NETs were defined as neutrophils positive for citrullinated histone 3.

### Spatial transcriptomic analysis of primary colon tumors and CRLM

Archived slides and scans from primary colon tumors and hard-set coverslips were removed by immersing in xylene overnight until coverslips were completely detached from the slide. After the coverslips were removed, slides were prepared and RNA was extracted according to the demonstrated protocol CG000518 (10x Genomics) using the Qiagen RNeasy FFPE Kit (Qiagen, PN:73504). RNA integrity was checked by the DV200 assay on the Agilent TapeStation with an RNA High Sensitivity kit (Agilent, PN:5067-5579) to assess the percent of RNA fragments larger than 200 nucleotides. Slides with DV200s of 50% or greater were used in the Visium assay. The Visium Human 6.5mm FFPE CytAssist assay (10x Genomics, PN:1000520) was performed according to manufacturer’s instructions (10x Genomics CG000495) and libraries were sequenced on a NextSeq 2000 to target 100 million reads per capture area. Sequenced reads were mapped to the Visium Human Transcriptome Probe Set v2.0 and aligned to their spots in the paired H&E images using Space Ranger 3.0.0 (10x Genomics).

Two 10X Visium ST colon cancer samples were analyzed using Seurat v5. The samples were preprocessed to include only spots within tissues and then underwent normalization and variance stabilization with sctransform in Seurat. Principal component analysis (PCA) was performed on the spots-features matrices with 50 components. Only 30 of the PCs were used as input for the FindNeighbors (k = 15) and UMAP functions in Seurat v5. With these parameters, we identified 22 clusters (resolution = 1.8) in the first sample and 28 clusters (resolution = 1.8) in the second sample. Differential gene expression (DGE) analysis was conducted for necrosis, peri-necrotic, and non-necrotic tumors using the FindMarkers function in Seurat v5. The genes that were positively regulated for each identity were selected. The adjusted p-value, calculated using Bonferroni correction, of 0.05 was used as cutoff to determine significance for regulated genes across these regions. The log1p- transformed feature counts were aggregated across the three different regions and then scaled before plotting the heatmaps.

Using the Wilcoxon Rank Sum test, we identified the positively upregulated gene set in the peri- necrotic areas of the colon tissues compared to the non-peri-necrotic regions. Genes with an adjusted p-value, calculated using Bonferroni correction, of less than 0.05 were used for pathway analysis. The gene sets from the two tissues were combined for a joint analysis of enriched pathways in GO and Reactome databases. Only terms with an adjusted p-value, calculated using Benjamini–Hochberg correction, of less than 0.05 were considered significant for the given gene set.

### Plasma cytokine analysis

Blood from humans or mice was collected in EDTA-coated tubes. Blood was centrifuged twice, first for 15 minutes at 1000g to obtain platelet rich plasma. Supernatant was collected and spun for 10 minutes at 10000g to obtain platelet poor plasma. Samples were stored at 80°C until use. Mouse Luminex® Discovery Assay and Human Luminex® Discovery Assay were used for mouse and human plasma, respectively. Samples and plates were prepared per manufacturer’s instructions and run by the Johns Hopkins Flow Cytometry Core Facility.

### Single cell RNA sequencing

Resected tumors were stored in DMEM with antibiotics and processed within 2-4 hours. First, the tumors were cut into ∼2 mm pieces, washed 3 times with DPBS, processed using tumor dissociation kit (Miltenyi Biotec), which uses mechanical and enzymatic digestion. Tumor fractions were incubated with dissociation media (DMEM and enzymes) and mechanically processed using a tissue dissociator (Miltenyi Biotec) at recommended soft tissue settings. Barcoded 3’ single cell libraries were prepared from the single cell suspensions using the Chromium Next GEM Single Cell 3’ kit v3.1(10X Genomics, Pleasanton, California) for sequencing according to the manufacturer’s instructions. The cDNA and library fragment size distribution were verified on a Bioanalyzer 2100 (Agilent, Santa Clara, CA). The libraries were quantified by fluorometry on a Qubit instrument (LifeTechnologies, Carlsbad, CA) and by qPCR with a Kapa Library Quant kit (Kapa Biosystems-Roche) prior to sequencing. The libraries were sequenced in a 28 x 10 x 10 x 90 bp configuration on an AVITI sequencer (Element Biosciences, San Diego, CA). The sequencing generated approximately 40,000 reads per cell. Data was then further analyzed using R. Low-quality cells were filtered if they have library sizes below 100,000 reads; express fewer than 2,000 genes; have spike-in proportions above 10%; or have mitochondrial proportions above 10%. Cell type annotation was performed using the Human Primary Cell Atlas reference using the SingleR (2.4.1) library. Data was harmonized using the Harmony library (1.2.0) and batch corrected with the Batchelor library (1.18.1). Then only cells identified as “Epithelial cells” were kept for subsequent analyses. The differentially expressed genes (DEGs) from the k-means cluster 1 vs. rest of clusters were calculated using DESeq2 (1.42.1), and pathway enrichment of those DEGs were calculated with gProfiler2 (0.2.3).

### RNA scope in-situ RNA hybridization

RNAScope in-situ RNA hybridization was performed using reagents, kits, and ACDBio protocol RNAscope Multiplex Fluorescent Reagent v2 (Advanced Cell Diagnostics, Document #: UM 323100). For FFPE tissue sections, tissue was cut to 5-μm thickness. Tissue was deparaffinized by placement in HybEZ™ II Oven at 60C for 1 hour followed by a histoclear series. Slides were then dehydrated in an ethanol series followed by placement in the oven at 60C for 5 minutes. Tissue sections were then treated with RNAScope hydrogen peroxide to cover and incubated for 10 minutes at room temperature. Slides were then prewarmed for 10 seconds in water maintained at boiling temperature then immediately transferred and incubated in 1X Target Retrieval Buffer maintained at a boiling temperature using a food steamer for 15 minutes. Slides were then rinsed in deionized water followed by dehydration using 100% ethanol and drying in the oven at 60C for 5 minutes. A hydrophobic barrier was drawn around tissue sections and allowed to dry. Slides were then treated with RNAScope Protease Plus to cover and incubated in the oven at 40C for 30 minutes. Sections were then washed in deionized water and subsequently incubated with probes (Hs-CDH-1, Hs-Snai1, Hs- Vimentin) at 40C for 2 hours. Slides were then washed overnight in prepared 5xSSC (1× SSC is 0.15 mol/L NaCl, 0.015 mol/L Na-citrate), at room temperature. Sections were then washed, and amplification steps were carried out using RNAScope Multiplex FL v2 Amp 1-3 reagents per kit. HRP signals were developed per protocol with channels and fluorophore corresponding to: CDH1 (Cy3-570); Snai-1 (Cy5-650); and Vimentin (FITC-520). Slides were then counterstained with DAPI and mounted with Prolong Gold Antifade then submitted for imaging.

### Chemotaxis assay

Transwells (Corning, # 3421) were pre-incubated with 200uL Chemotaxis Media (RPMI + 0.5% BSA) for 30 minutes at 37°C. Blood from healthy volunteers was collected and lysed using ACK buffer. Cells were washed with twice with CM. 100uL of the cell solution was added to the transwells. 600uL of CM was added to the wells with (treated) or without (untreated) 5uL/mL of CXCL1 (R&D, #9196- GR-025/CF). Control wells of 100uL cell solution in 500uL CM was prepared. Plates were incubated and cells were allowed to migrate for 3 hours at 37°C. Following incubation, transwells were carefully removed. 500uL of solution from the treatment wells and control wells were collected and washed. Solutions were stained and analyzed as previously stated (see “*Flow cytometry and cell sorting*”).

### In vivo human neutrophil migration assay

Healthy volunteers donated blood and then washed their mouth three times with 15mL of sterile saline for 30 seconds. Washout was recovered. Volunteers then washed their mouths for 30 seconds with 10% Tabasco solution in saline and were asked to avoid eating and drinking. Two hours later, they again washed their mouths three times with sterile saline, and washout was recovered. All mouthwash and blood samples were centrifuged, stained, and analyzed as previously stated (see “*Flow cytometry and cell sorting*”).

### Statistical analysis

Unless otherwise indicated, data are represented as mean values ± standard error of the mean (SEM). Paired or unpaired Student’s t-tests were used to compare two groups, and more than two datasets were compared using one-way analysis of variance (ANOVA) with Tukey’s post-test. Where applicable, normality was estimated using D’Agostino & Pearson or Shapiro-Wilk normality tests. Log-rank (Mantel-Cox) analysis was used for Kaplan-Meier survival curves. The tests used are stated in the Figure legends. No samples were excluded. All statistical analyses, except for RNA-seq analyses, InfinityFlow, and spatial transcriptomics (see corresponding sections above), were performed using Prism v9 (GraphPad, California, USA). A p-value < 0.05 was considered statistically significant; non-significant (n.s.) differences are indicated in the Figures.

### Data and code availability

Human CRC liver metastasis scRNAseq data is available at GEO, accession number: GSE283816. Spatial transcriptomics data is available at GEO, accession number: GSE281023. All other pieces of data are available upon request.

## Statement of significance

Neutrophil extracellular traps (NETs) induce colorectal tumor necrosis and drive cancer cell metastasis. Pharmacological targeting of NETs decreases tumor necrosis and metastasis, and enhances the efficacy of treatment with standard of care chemotherapy.

