## Supplementary data for "Targeting Neutrophil Extracellular Traps to inhibit Colon Cancer Tumor Necrosis and Metastasis"

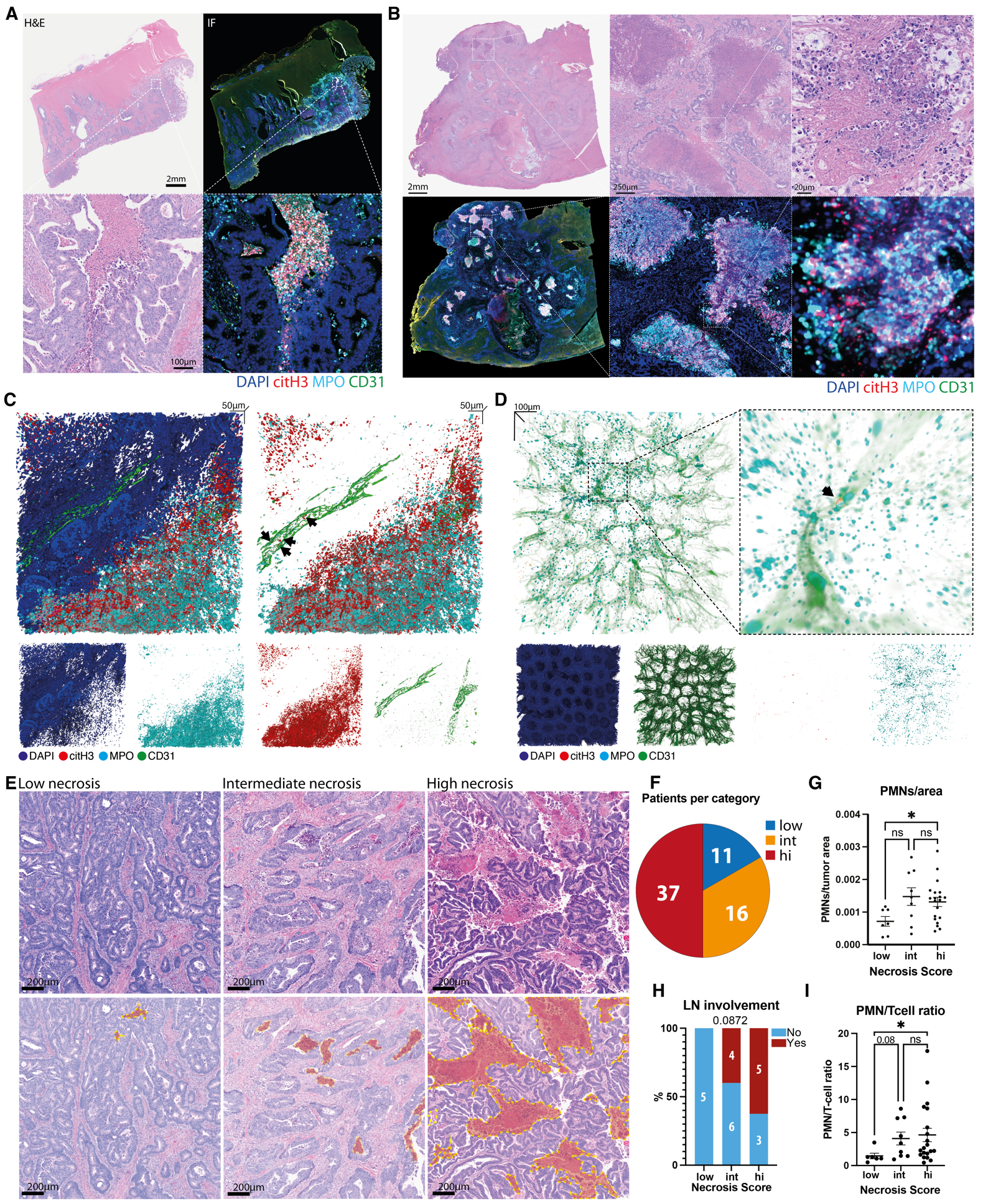

**Supplementary Figure 1.** **Necrosis is associated with NET accumulation and vascular occlusion in patients with colon cancer** **A)** Representative H&E-stained and immunostained human colon tumors and **B)** liver metastases, showing necrosis and NETs. Representative of N=45 colon tumors and N=16 liver metastases. **C)** Representative image of whole mount cleared liver metastasis showing disruption of intra-tumoral vasculature in necrotic regions, and NETs (black arrows) inside intact vessels outside of necrotic areas. Representative of N=4 liver metastases. **D)** Representative image of whole mount cleared adjacent normal colon from a patient with colon cancer showing presence of NETs. Representative of N=4 normal colon tissues. **E)** Examples of tumors showing low, intermediate and high necrosis score. **F)** Distribution of patients into groups based on necrosis score. **G)** Quantification of neutrophils per tumor area, **H)** locoregional lymph node invasion and **I)** neutrophil to T-cell ratio in low, intermediate and high necrotic CRC tumors. Graphs show mean ± s.e.m., *P< 0.05, **P< 0.01, ***P< 0.001, n.s., not significant, as determined by Fisher’s exact test (H), or one-way ANOVA with Dunnett’s multiple comparison test (G, I).

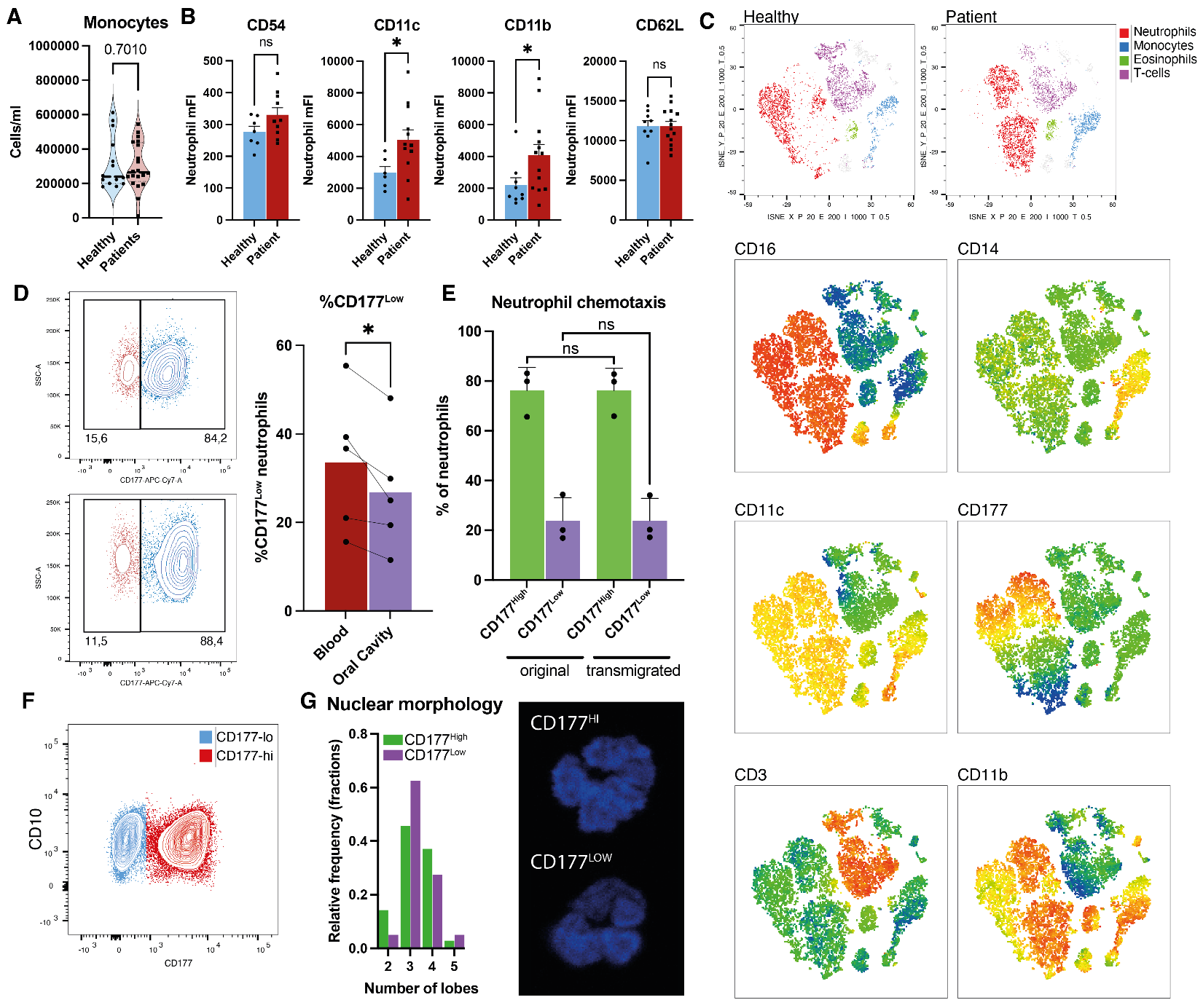

**Supplementary Figure 2. Altered neutrophil populations and reduced extravascular capacity of NET-forming neutrophils in CRC patients A)** Quantification of monocytes in circulation of patients and healthy controls. N=20 patients and N=14 controls. **B)** Quantification of different makers in circulating neutrophils (DAPI^-^, CD45^+^, CD16^+^) from healthy controls and CRC patients. **C)** Representative low dimensional representation of blood immune populations (top) and marker abundances (bottom). **D)** Representative plot and quantification of CD177^Hi^ and CD177^Low^ neutrophils in the blood prior to mucosal injury and in the oral cavity following mucosal stimulation, quantified by flow cytometry. Lines join data from the same participant **E)** Percent of CD177^Hi^ and CD177^Low^ neutrophils before and after chemotaxis in a transwell assay. N= 3. **F)** Representative plot of CD10 expression in CD177^HI^ and CD177^Low^ neutrophils, as quantified by flow cytometry. **G)** Histogram (left) and representative images (right) of nuclear morphology of CD177^Hi^ and CD177^Low^ neutrophils. Bars show mean ± s.e.m., *P< 0.05, n.s., not significant, as determined by one-way ANOVA with Dunnett’s multiple comparison test (E), paired two-tailed t-test (D) or unpaired two-tailed t-test (A, B).

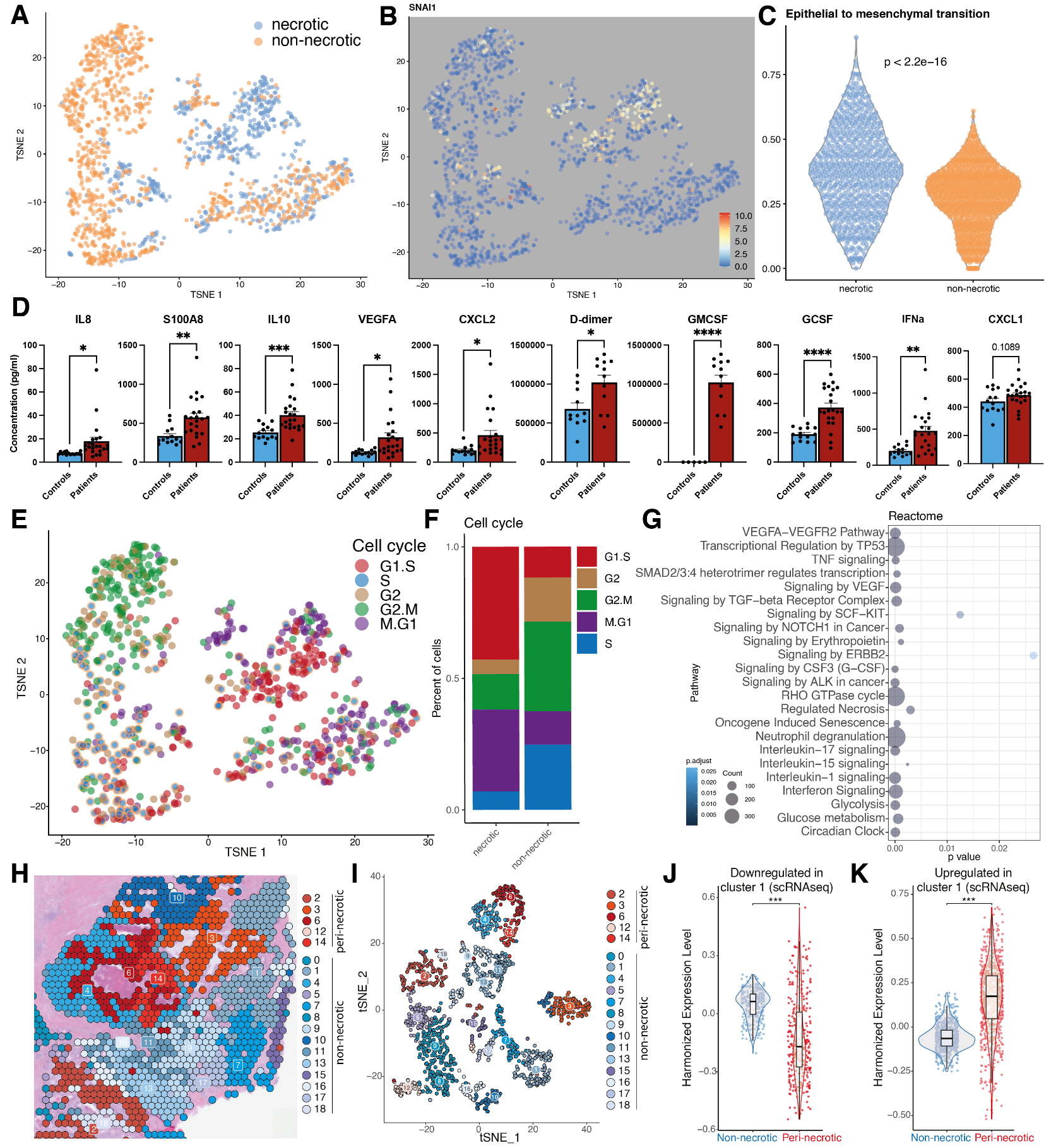

**Supplementary Figure 3.**  **Necrotic Tumor Microenvironments Reprogram Epithelial Cells Toward EMT and Pro-Metastatic Pathways A)** UMAP of epithelial cells from necrotic and non-necrotic tumors. N=2 necrotic and N=2 non-necrotic tumors **B)** Expression of SNAI1 in epithelial cells of necrotic and non-necrotic tumors. **C)** Aggregated score of epithelial-to-mesenchymal pathway in epithelial cells of necrotic and non-necrotic tumors. P-value from Wilcoxon test. **D)** Quantification of the levels of several chemokines, cytokines and colony stimulating factors in the plasma of CRC patients (n=22) compared to healthy volunteers (n=14). **E)** UMAP and **F)** quantification of the percent of epithelial cells from necrotic and non-necrotic tumors in the different cell cycle phases. **G)** Analysis of Reactome pathways enriched in epithelial cells of necrotic compared to non-necrotic tumors. **H)** Overlay of the detected clusters and **I)** UMAP of the unbiased clustering of our spatial transcriptomic dataset. **J-K)** Aggregated expression of the downregulated (**J**) and upregulated (**K**) genes in cluster 1 of the single cell sequencing in peri-necrotic and non-peri-necrotic tumor regions of our spatial transcriptomics dataset.

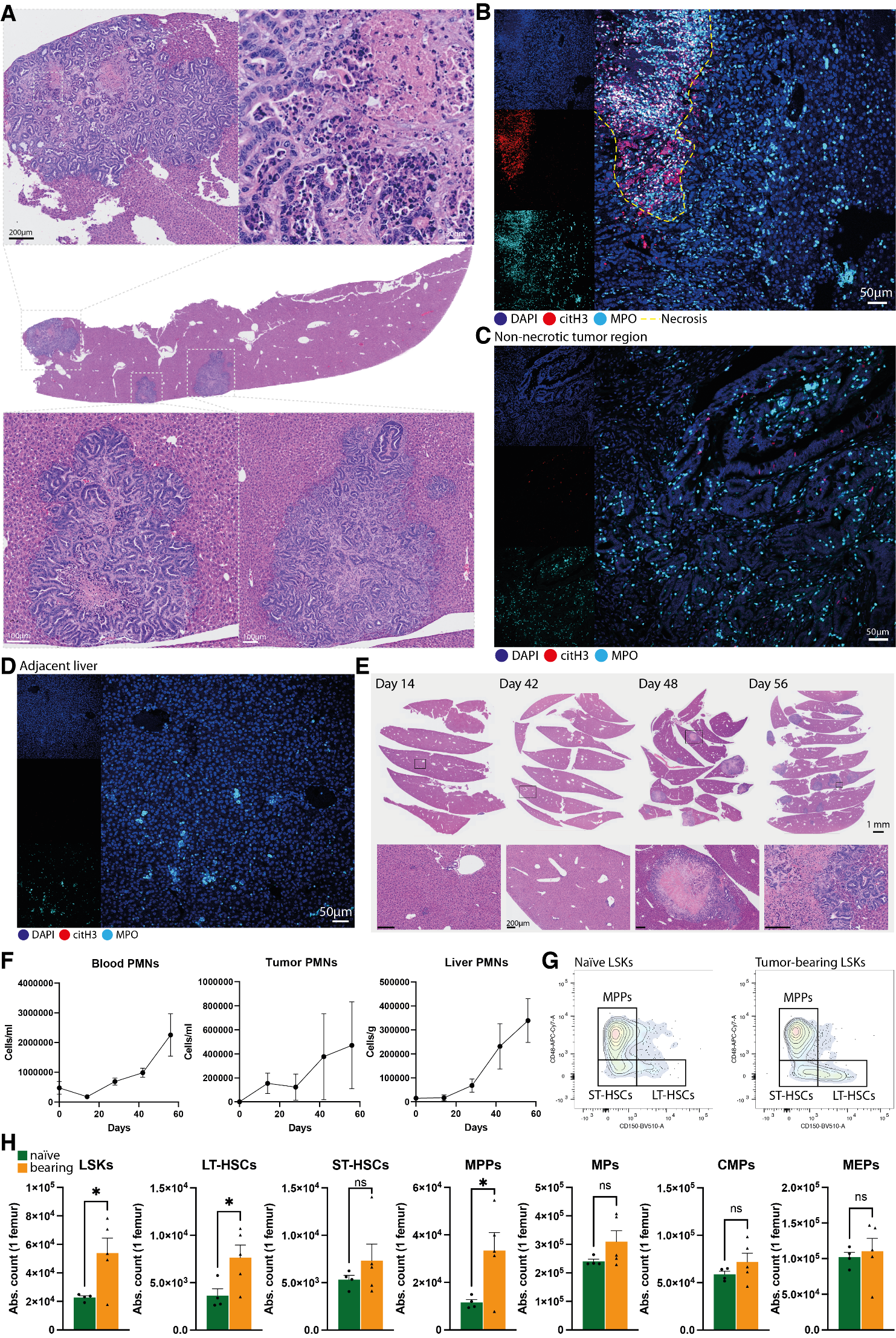

**Supplementary Figure 4. Necrosis-associated NET accumulation in liver metastases and myeloid expansion in a CRC A)** Representative image of H&E-stained liver metastases with varying levels of necrosis in mice with AKPS colon tumors. Representative of N=6 livers. **B)** Representative immunostaining of NETs in non-necrotic regions, **C)** necrotic regions of metastases, and **D)** in adjacent normal liver tissue in mice with AKPS colon tumors. Representative of N=3 livers. **E)** H&E-stained images of liver during AKPS colon tumor progression. Representative of 3 livers per timepoint. **F)** Number of neutrophils in circulation, within colon tumors, and within livers during tumor progression, quantified by flow cytometry. N=3 mice per timepoint. **G)** Representative cytometry plots of lineage-negative cells in the bone marrow of tumor-bearing and naïve mice. N=5 tumor-bearing mice and N=4 naïve mice. **H)** Number of cell populations in the bone marrow of tumor-bearing and naïve mice. N=5 tumor-bearing mice and N=4 naïve mice. LSKs: Lineage^–^, Sca1^+^, cKit^+^; LTC-HSCs: long-term hematopoietic stem cells; ST-HSCs: short-term hematopoietic stem cells; MPP: multipotent progenitors; MP: myeloid progenitors; CMPs: common myeloid progenitors; MEPs: megakaryocyte erythrocyte progenitors. Bars show mean ± s.e.m., *P< 0.05, n.s., not significant, as determined by unpaired two-tailed t-test.

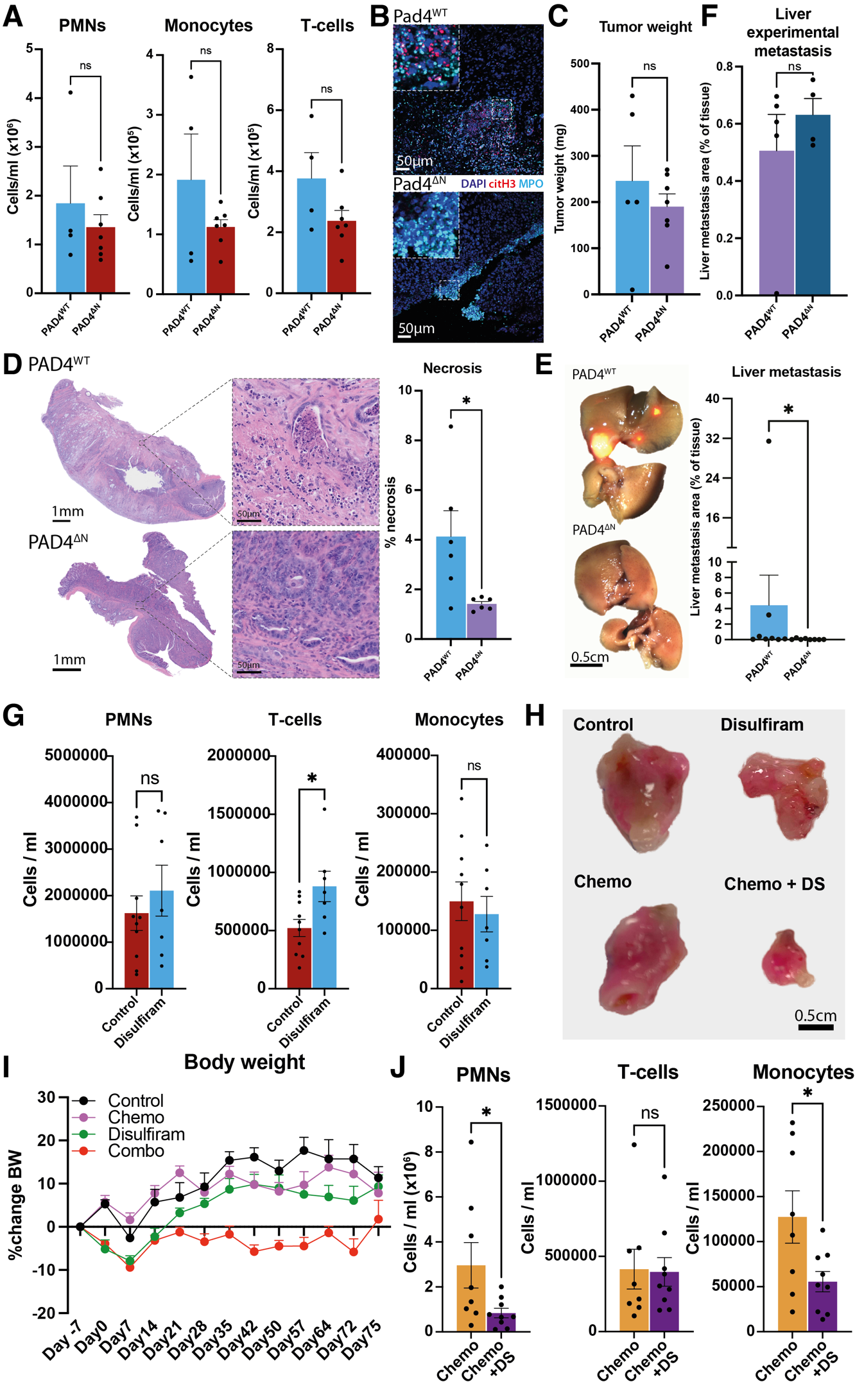

**Supplementary Figure 5. PAD4 deficiency decreases tumor necrosis and metastatic spread in CRC models A)** Number of circulating neutrophils, monocytes, and T-cells of PAD4^KO^ and PAD^ΔN^ mice, as quantified by flow cytometry. N=4 PAD4^WT^ and N=7 PAD4^ΔN^ mice**. B)** Representative image of immunostained NETs in colon tumors of PAD4^WT^ and PAD4^ΔN^ mice. N=5 PAD4^WT^ and N=7 PAD4^ΔN^ mice. **C)** Quantification of colon tumor weight of PAD4^WT^ and PAD4^ΔN^ mice at endpoint. N=5 PAD4^WT^ and N=7 PAD4^ΔN^ mice. **D)** Representative H&E-stained images and quantification of necrosis within the colon tumors of PAD4^WT^ and PAD^ΔN^ mice. N=6 PAD4^WT^ mice and N=6 PAD^ΔN^ mice. **E)** Representative brightfield images with overlying tumoral mScarlet fluorescence and quantification of liver metastasis of PAD4^WT^ and PAD^ΔN^ mice. N=8 PAD4^WT^ mice and N=8 PAD^ΔN^ mice. **F)** Metastatic area in the livers from AKPS-tumor bearing PAD4^WT^ and PAD4^ΔN^ mice subjected to experimental metastasis through the portal vein. N=5 PAD4^WT^ and N=4 PAD4^ΔN^ mice. **G)** Number of circulating neutrophils, monocytes, and T cells of control and disulfiram-treated mice, as quantified by flow cytometry. N=10 control mice and N=7 disulfiram-treated mice. **H)** Representative images of AKPS colon tumors of each treatment group at endpoint. **I)** Percent change in bodyweight of tumor-bearing mice throughout treatment. **J)** Number of circulating neutrophils, monocytes, and T-cells of chemotherapy-treated and combination-treated mice, as quantified by flow cytometry. N=8 chemotherapy treated and N=9 combination-treated mice. Bars show mean ± s.e.m., *P< 0.05, n.s., not significant, as determined by unpaired two-tailed t-test.

### Supplementary video legends

**Supplementary Video 1:** A primary colon tumor from a CRC patient cleared and stained for the vasculature (CD31, green), neutrophils (MPO, cyan), citrullinated histone 3 (as a marker of NETs, red), and DAPI (blue), showing necrotic regions enriched in neutrophils and NETs and devoid of intact vasculature.

**Supplementary Video 2:** A primary colon tumor from a CRC patient cleared and stained for the vasculature (CD31, green), neutrophils (MPO, cyan), citrullinated histone 3 (as a marker of NETs, red), and DAPI (blue), showing necrotic regions enriched in neutrophils and NETs and devoid of intact vasculature, and presence of NETs in vessels in regions surrounding necrosis.

**Supplementary Video 3:** A primary colon tumor from a CRC patient cleared and stained for the vasculature (CD31, green), neutrophils (MPO, cyan), citrullinated histone 3 (as a marker of NETs, red), and DAPI (blue), showing a reconstruction of a vessel leading into a necrotic region, and its intraluminal contents showing abundant neutrophil and NETs aggregates.

**Supplementary Video 4:** A normal adjacent colon from a CRC patient cleared and stained for the vasculature (CD31, green), neutrophils (MPO, cyan), citrullinated histone 3 (as a marker of NETs, red), and DAPI (blue), showing presence of NETs in extra-tumoral vessels.
